# Nanoporous Platinum Microelectrode Arrays for Neuroscience Applications

**DOI:** 10.1101/2024.06.06.597813

**Authors:** Nicolai Winter-Hjelm, Leik Isdal, Peter A. Köllensperger, Axel Sandvig, Ioanna Sandvig, Pawel Sikorski

## Abstract

Microelectrode arrays are invaluable tools for investigating the electrophysiological behaviour of neuronal networks with high spatiotemporal precision. In recent years, it has become increasingly common to functionalize such electrodes with highly porous platinum to increase their effective surface area, and hence their signal-to-noise ratio. Although such functionalization significantly improves the electrochemical performance of the electrodes, the impact of various electrode morphologies on biocompatibility and electrophysiological performance in cell cultures remains poorly understood. In this study, we introduce reproducible protocols for depositing highly porous platinum with varying morphologies on microelectrodes designed for neural cell cultures. We also evaluate the impact of morphology and electrode size on the signal-to-noise ratio in recordings from rat cortical neurons cultured on these electrodes. Our results indicate that electrodes with a uniform layer of highly nanoporous platinum offer the best trade-off between biocompatibility, electrochemical, and electrophysiological performance. While more microporous electrodes exhibited lower impedance, nanoporous electrodes detected higher extracellular signal amplitudes from neurons, suggesting reduced distance between perisomatic neuronal areas and the electrodes. Additionally, these nanoporous electrodes showed fewer thickness variations at their edges compared to the more porous electrodes. Such edges can be mechanically broken off during cell culturing and contribute to long-term cytotoxic effects, which is highly undesirable. We hope this work will contribute to better standardization in creating and utilizing nanoporous platinum microelectrodes for neuroscience applications. Improving the accessibility and reproducibility of this technology is crucial for enhancing the quality of electrophysiological data and advancing our understanding of neuronal network function and dysfunction.

## Introduction

Microelectrode arrays (MEAs) are extensively used to study neuronal signaling and behaviour from the cellular to the network level (1–3). These microelectrodes detect voltage fluctuations caused by propagating action potentials, commonly referred to as spikes, with high temporal precision. By monitoring the spiking activity of tens to thousands of neurons simultaneously, key principles of neuronal function and dysfunction can be deciphered (4, 5). Optimizing MEAs for high-quality neuronal recordings typically involves a trade-off between high selectivity and a high signal-to-noise ratio (SNR). Ideally, electrodes should be the size of individual neurons, ranging from 10 to 100 µm, to ensure high selectivity. However, the electrode surface area is inversely proportional to impedance. To increase SNR without increasing electrode diameter, porous platinum has been employed (6–8), including the functionalization of microelectrodes for neuroscience applications (9–13). Porous platinum, also known as platinum black, is highly permeable platinum that absorbs a large portion of incoming light, appearing black under a microscope (14, 15). Its high porosity creates a large electrochemically active surface area, significantly reducing electrode impedance. The increased surface area additionally generates higher currents in response to applied potentials, allowing for lower voltages to stimulate neuronal networks. Consequently, the likelihood of exceeding the so-called water window is reduced. Exceeding this threshold can trigger oxidation and reduction reactions, leading to electrolysis, which can be detrimental to nearby biological tissues (10). Thus, porous platinum can substantially enhance both the electrophysiological performance and biocompatibility of extracellular microelectrodes.

Platinum black can be formed through various fabrication methods, ranging from bottom-up electrochemical synthesis to laser roughening (16, 17). These methods yield surfaces with a range of morphologies (18, 19). One of the most reliable methods for forming platinum black is electrochemical deposition, also know as electroplating (9, 10, 20, 21). In this process, electrodes are submerged in a liquid containing platinum ions, typically chloroplatinic acid (22). An external potential is applied to electrochemically reduce the platinum ions, which then form nuclei and grow into dendritic protrusions. Traditionally, additives such as lead have been introduced to the bath. While the exact nucleation and growth mechanisms of platinum nano- and microstructures are not fully understood, various additives are believed to either inhibit the growth of platinum islands (23) or assist in electron transfer between the electrode and the platinum ions (24, 25). However, these additives can leak into cell media and induce cytotoxicity, and they complicate the deposition process due to the complexity of the electrochemical reactions involved (12). In 2015, Boehler *et al*. introduced an alternative process using formic acid in the electrolyte solution, proposed to have less impact on cell viability (10). Later, they demonstrated that depositions could occur even without any additives (12).

A challenge in fabricating porous microelectrodes is the unpredictability of the deposition process, which can result in heterogeneous metallic protrusions across the surface. Many publications report using platinum black electrodes without characterizing their morphology or biocompatibility. The deposition kinetics often lead to the formation of high, irregular edges, commonly referred to as edge effects, that can impede neuronal growth and increase the neuron-electrode junction size. While signals can still be acquired from neurons near the electrodes, these edges prevent neurons from growing directly on top of the electrodes. Additionally, these metallic protrusions can break off during fabrication or use with cell cultures. Although platinum is widely used in biological and medical electrode applications due to its inertness and high electrical conductivity (26), studies have shown that platinum nanoparticles can cause cytotoxic and inflammatory effects in neurons both *ex vivo* and *in vivo* (27, 28). Therefore, it is desirable to have uniformly layered platinum deposits without discernible protrusions that could break off. Wang *et al*. demonstrated that such electrodes could be achieved by applying constant currents to the electrodeposition system (13). However, using constant current for electroplating presents challenges, as the same deposition parameters cannot be reliably used for different electrode sizes or heights (i.e., deposition times). This is because the depositions continuously change the surface area and thus the current distribution in the system. This variability makes the depositions highly inconsistent, as even minor changes in electrode conditions can lead to significant variations in the morphologies of the platinum deposits. Therefore, more reproducible and standardized protocols are needed for creating nanoporous microelectrodes for neuroscience applications.

In this study, we present a reproducible method for fabricating nanoporous, biocompatible microelectrodes with uniform thickness for bioelectronic applications. We compare various electrochemical deposition schemes to demonstrate their impact on the nucleation and growth of nanoporous platinum and evaluate their electrochemical performance. Additionally, we investigate how the resulting structures affect electrophysiological recordings from cultured neurons. Ultimately, we propose an optimal deposition scheme that balances biocompatibility, electrochemical performance, and electrophysiological performance for use in neuronal recordings.

## Materials and Methods

### Design of Experimental Interfaces

Designs for all utilized interfaces were created using Clewin 4 (WieWeb Software, Enschede), and are shown in **Figure S1**. To optimize the electrochemical deposition parameters and characterization, 20 electrodes were connected to individual contact pads along the periphery of a 4” wafer (**Figure S1A**). Electrodes with diameters ranging from 10 µm to 100 µm were used for optimizing deposition parameters, while all electrochemical characterizations were conducted using 30 µm diameter electrodes. The metal layer was made 5 µm wider than the etching layer to increase alignment tolerance. For electrodes to be tested with neural cell cultures, designs were made compatible with a MEA2100 workstation from Multichannel Systems. These platforms (n = 6) consisted of 59 electrodes with 30 µm diameter and 600 µm interspace (**Figure S1B**). 1/3 of the electrodes acted as planar controls, 1/3 were electroplated with nanoporous platinum at −0.4 V, and 1/3 were electroplated with microporous platinum at −0.3 V. Each electrode was connected to a contact pad compatible with the recording hardware. Larger contact pads positioned at the periphery of the wafer connected up to 15 electrodes each to the potentiostat during electroplating. These connections were broken once the wafers were diced into 49 x 49 mm substrates compatible with the MEA2100 workstation. 6.0 mm diameter PDMS chambers were used as cell compartments. Samples for scanning electron microscopy were created with 520 planar control electrodes, 520 electrodes electroplated with platinum at −0.4 V, and 520 electrodes electroplated at −0.3 V (**Figure S1C**).

### Fabrication of Microelectrode Arrays

The protocol for fabricating MEAs was adapted from our previous work (29, 30) and modified to support electrochemical depositions of platinum. An illustration of the fabrication steps is shown in **Figure 1A**.

**Figure 1.**
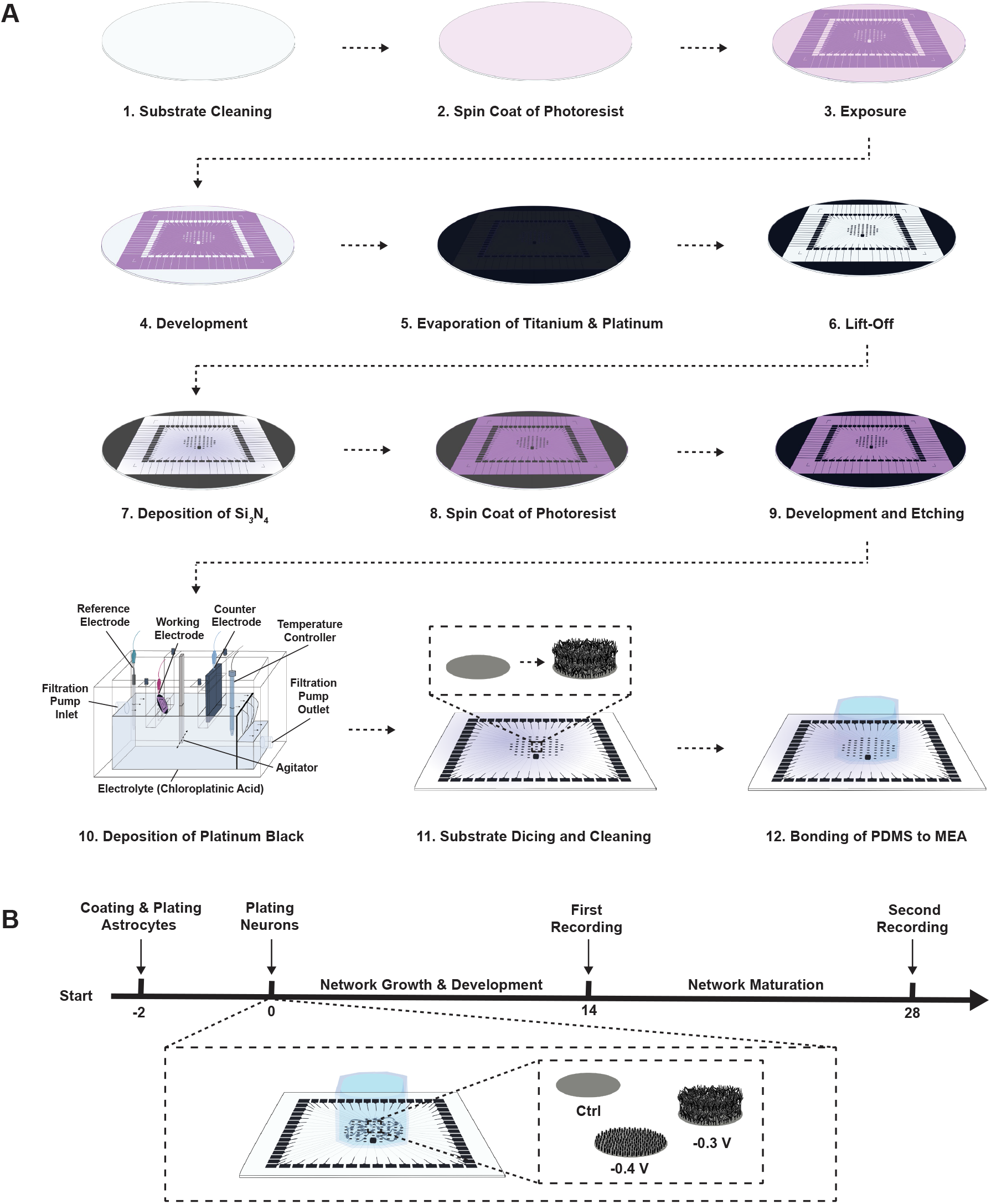
Experimental Setup. **(A.)** Illustration of the fabrication procedure for creating microelectrode arrays with electroplated porous platinum. **(B.)** Experimental timeline for cell experiments with co-cultures of primary rat cortical neurons and astrocytes. Each utilized platform had 1/3 planar control electrodes, 1/3 highly microporous, heterogeneous electrodes electroplated at −0.3 V, and 1/3 highly nanoporous, evenly layered electrodes electroplated at −0.4 V. Electrophysiological recordings were conducted at 14 and 28 *days in vitro* (DIV) for 15 min each.

MEAs were fabricated on 1 mm thick 4-inch borosilicate wafers (100 mm Borofloat33, Plan Optik). First, the wafers were washed in acetone and IPA for 1 min each to remove organic contaminants, followed by a 5 min plasma clean in 100 sccm O_2_ plasma at 20 kHz generator frequency (Femto Plasma Cleaner, Diener Electronics). The wafers were then dehydration baked at 100 °C for 2 min to remove moisture. A 4 µm thick layer of the photoresist ma-N 440 (micro resist Technology GmbH) was spin coated onto the substrates at 3000 rpm for 42 s with a ramp rate of 500 rpm*/*min (spin150, SPS-Europe B.V.). The films were left to rest for 10 min before a soft bake was conducted at 95 °C for 5 min. For exposures of the design, a maskless aligner (MLA150, Heidelberg) with a 405 nm laser at 1800 mJ*/*cm^2^ was used. Development was conducted in ma-D332/s (micro resist Technology GmbH) for 90 ± 10 s, followed by thorough rinsing in deionized (DI) water. Next, a short descum was performed using 100 sccm O_2_ plasma for 1 min at 20 kHz generator frequency. A 50 nm titanium adhesion layer was then evaporated onto the substrates at 5 Ås^−1^, followed by 100 nm platinum at 2 Ås^−1^ (E-beam Evaporator, Pfeiffer Vacuum Classic 500). Finally, a lift-off step was conducted in acetone, followed by a thorough rinse in IPA. Another 1 min descum was conducted prior to the deposition of a passivation layer. Subsequently, 470 nm silicon nitride (Si_3_N_4_) was deposited on the substrates using plasma-enhanced chemical vapour deposition (PECVD) at 300 °C for 30 min with 20.0 sccm SiH_4_, 20.0 sccm NH_3_, and 980 sccm N_2_ gas (PlasmaLab System 100-PECVD, Oxford Instruments). Thereafter, an etch mask was created using a second layer of ma-N 440, following the same protocol as previously described. Finally, the silicon nitride above the electrodes and contact pads was dry etched using inductively coupled plasma (ICP) with 50.0 sccm CHF_3_, 10.0 sccm CF_4_, and 7.0 sccm O_2_ gas for 6.5 min (Plasmalab System 100 ICP-RIE 180, Oxford Instruments).

### Electroplating of Porous Platinum

A PP-Type Wafer Plating Laboratory System from Yamamoto was used for electrodepositions. The electrolyte solution consisted of aqueous 2.5 mmol chloroplatinic acid (H_2_PtCl_6_, 8 wt% H_2_O, Sigma-Aldrich, 262587). In some experiments, 1.5 mmol formic acid (HCOOH, 95%, Sigma-Aldrich, W248703) was added to the bath as an additive. A custom-built wafer holder was used to position the wafer in the electrolyte bath so that the liquid covered the working electrodes but not the contact pads (**Figure 1A**). A Red Rod REF201 Ag/AgCl electrode (Hatch) was used as the reference electrode, and a platinized titanium plate served as the counter electrode. An external temperature controller maintained the temperature at 30 °C. A paddle agitator, placed between the working electrode and the counter electrode, was kept at 60.0 rpm to reduce the diffusion layer thickness. A PalmSens 4 potentiostat (PalmSens) was used in various modes, including chronoamperometric, chronopotentiometric, linear sweep voltammetry, and cyclic voltammetry, for all depositions.

### Preparation of PDMS Compartments for Cell Experiments

To prepare PDMS compartments for the cell cultures, silicon elastomer and curing agent (SYLGARD®184 elastomer kit, Dow Corning) were mixed, degassed, and cast at a ratio of 10:1 on top of a precleaned 4-inch silicon wafer (Siltronix). The PDMS was then cured at 65 °C for 4 h (TS8056, Termaks). After curing, the PDMS was peeled from the mold, and a 6 mm diameter punch was used to cut out the cell compartments. Debris was removed from the PDMS chips using scotch tape, and the chips were washed successively in acetone, 96% ethanol, and DI water for 1 min each. Prior to bonding, the chips were left to dry overnight in a fume hood.

### Assembly of Microdevices

The MEAs were cut into 49 × 49 mm substrates compatible with the Multichannel head stage using a wafer saw (DAD323, DISCO). Following this, the ma-N 440 layer was removed using acetone and IPA. A 10 min plasma clean was conducted using 160 sccm O_2_ at 32 kHz generator frequency to remove hardened photoresist and oxidize the top nanometers of the silicon nitride layer into silicon dioxide (SiO_2_). To activate the surfaces of the PDMS device and MEA for bonding, an O_2_ plasma treatment was conducted using 200 sccm O_2_ plasma at 40 kHz generator frequency for 1 min. Subsequently, two drops of 70% ethanol were placed between the MEA and the PDMS chip, facilitating alignment of the PDMS compartment above the microelectrodes under a stereomicroscope. The bonding was completed on a hotplate at 100 °C for 90 s, followed by 5 min at room temperature under gentle pressure. DI water was used to wash out any remaining traces of ethanol and to keep the surfaces of the MEAs hydrophilic. Prior to cell culturing, the devices were sterilized under UV light in a biosafety cabinet for at least 60 min.

### Electrochemical Characterization

As the electrolyte, 0.1 M Dulbecco’s Phosphate Buffered Saline (DPBS) (Sigma-Aldrich, D8537) was used. A custom-built wafer holder was employed to position the wafer in the electrolyte bath, ensuring the liquid covered the working electrodes but not the contact pads. A Red Rod REF201 Ag/AgCl electrode (Hatch) was used as the reference electrode, and a platinized titanium plate served as the counter electrode. A PalmSens 4 potentiostat (PalmSens) was used to perform electrochemical impedance spectroscopy (EIS) measurements. Nine electrodes of each condition were analyzed, and the average curve of the nine electrodes was plotted. An equilibration time of 8 ms was set. Measurements were performed against the open circuit potential (OCP), with the constant potential (E dc) set to 0 V. An alternating sine wave of 100 mV was superimposed on the constant potential. This amplitude was set high to avoid issues with noise at low frequencies and was confirmed not to impact the linearity of the system. The OCP was measured prior to the EIS scan with a maximum time of 3 s. A stability criterion of 0.5 mVs^−1^ was chosen. Impedance was measured in the range from 100,000 to 0.1 Hz with 10 measurement points per decade on a logarithmic scale. Data from the EIS measurements were validated using the Lin-KK method, based on the Kramers-Kronig (KK) compliant equivalent circuit (31).

### Coating, Cell Plating and Maintenance

An overview of the timeline for the cell experiments can be seen in **Figure 1B**. After sterilization and prior to coating the surfaces, the samples were soaked in DMEM, low glucose (Gibco™, 11885084) for at least 48 h to remove any potential uncured PDMS. The cell medium was then replaced with 0.1 mg*/*mL Poly-L-Ornithine (PLO) solution (Sigma-Aldrich, A-004-C), and the chips were incubated in a refrigerator at 4 °C overnight. The following day, the chambers were washed three times with MQ water before being filled with laminin solution consisting of 16 µg*/*mL natural mouse laminin (Gibco™, 23017015) in phosphate-buffered saline (PBS, Sigma-Aldrich, D8537). The samples were then incubated at 37 °C with 5% CO_2_ for 2 h.

Following coating, the laminin solution was replaced with pre-warmed astrocyte medium consisting of DMEM, low glucose, supplemented with 15% Fetal Bovine Serum (Sigma-Aldrich, F9665) and 2% Penicillin-Streptomycin (Sigma-Aldrich, P4333). Rat astrocytes (Gibco™, N7745100) were then plated at a density of 100 cells*/*mm^2^, equalling 4000 cells per culturing chamber. After two days of expansion, the astrocyte medium was replaced with neuronal medium consisting of Neurobasal Plus Medium (Gibco™, A3582801), supplemented with 2% B27 Plus (Gibco™, A358201), 1% GlutaMax (Gibco™, 35050038), and 2% Penicillin-Streptomycin (Sigma-Aldrich, P4333). Additionally, Rock Inhibitor (Y-27632 dihydrochloride, Y0503, Sigma-Aldrich) was added at a concentration of 0.1% during plating to increase the survival rate. Rat cortical neurons from Sprague Dawley rats (Gibco, A36511) were plated at a density of 1000 cells*/*mm^2^, equalling 40,000 cells per chamber. Half of the cell medium was replaced with fresh medium 4 h after plating, and again after 24 h. From then on, half of the cell medium was replaced every second day until the cultures were ended at 28 days *in vitro* (DIV). All astrocytes and neurons were from the same batch and cell vials to avoid biases caused by batch-to-batch variability.

### Electron Microscopy

To prepare biological samples for imaging, the cells were fixed using 2.5% glutaraldehyde (G5882, Sigma-Aldrich) in Sorensen’s phosphate buffer (0.1 M, pH 7.2) for at least 24 h. Initially, the cells were left in the fixative at room temperature for the first 2 h, followed by storage at 4 °C until chemical dehydration. Subsequently, a chemical serial dehydration was conducted using 25%, 50%, 70%, 80%, 90%, and 100% (x2) ethanol for 5 min each. The ethanol was then replaced by CO_2_ with critical point drying (EM CPD300 Critical Point Dryer, Leica). For this process, the delay time was set to 120 s, the exchange speed to 1, the number of cycles to 18, and the gas outflow speed to 50% (slow). To make the samples conductive, 15 nm of platinum/palladium was deposited on the samples using a 208 HR B Sputter Coater from Cressington. These depositions were conducted cyclically from −45^°^ to 45^°^ with a period of 20 s. All electron microscopy imaging was performed using an APREO Field Emission Scanning Electron Microscope from FEI. The samples were connected to the stage using copper tape. Imaging was conducted using secondary electrons detected with an EDT detector, with the beam current set between 25–50 pA, and the acceleration voltage between 4.0–10.0 kV.

### Electrophysiological Recordings

A MEA2100 workstation (Multichannel Systems) was used for recordings of neuronal activity, with the sampling rate set to 25000 Hz. Furthermore, the temperature was maintained at 37 °C using a temperature controller (TC01, Multichannel Systems). To maintain sterility during recordings, a 3D-printed plastic cap with a gas-permeable membrane was employed. Prior to recordings, the neuronal networks were allowed to equilibrate for 5 min at the recording stage. Subsequently, neuronal activity was recorded for 15 min. All recordings were conducted 24 h following media changes.

### Data Analysis and Statistical Analysis

Data analysis was performed using Matlab R2021b. To preprocess the data, a 4th order Butterworth bandpass filter was applied to eliminate frequencies below 300 Hz and above 3000 Hz. Additionally, noise originating from the power supply mains at 50 Hz was attenuated using a notch filter. These filters were implemented using zero-phase digital filtering to mitigate group delay in the output signal. For spike detection, the Precise Timing Spike Detection (PTSD) algorithm developed by Maccione *et al*. (32) was employed due to its demonstrated precision in detecting spikes from rat cortical and hippocampal neurons, validated through visual inspection. Unlike threshold-based methods, PTSD incorporates adaptive filtering and precise timing analysis, reducing false positives and ensuring consistent detection across varying electrode types. These strengths made it the most reliable choice for our study, allowing for accurate comparisons of neuronal activity and electrode performance. The data was thresholded at 9 times the standard deviation of the noise, with the maximum peak duration and refractory time set to 1 ms and 1.6 ms, respectively. This threshold was chosen to ensure a robust balance between sensitivity and specificity, capturing true spikes while avoiding the detection of minor fluctuations in the noise band as spikes. The threshold selection was validated through visual inspection of spike detection across a representative selection of electrodes, confirming that the detected events corresponded to clear neuronal spikes. The signal-to-noise ratio (SNR) was calculated in decibel (dB) using the formula:

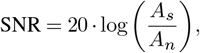

where *A*_*s*_ is the average spike amplitude recorded by the electrode, and *A*_*n*_ is the average amplitude of noise band fluctuations recorded by the same electrode.

Color maps for plotting were generated using the Matlab function *linspecer* (33), which is based on the web tool colorBrewer developed by Brewer *et al*. (34). All statistical analyses were conducted in Matlab R2021b using a two-sided Wilcoxon rank sum test.

## Results

### Applied Deposition Potential Determines the Electrode Porosity

To evaluate the impact of the applied deposition potential on the porosity of platinum microelectrodes, chronoamperometric depositions were conducted in the range from 0 V to − 0.4 V. The porosity increased with reduced deposition potential down to −0.3 V, as indicated by the darker color and more discernible thickness variations at the electrode periphery when viewed under a microscope (**Figure 2A**). While the amount of deposited platinum was highest at − 0.4 V, the light absorbance was less pronounced, indicating lower porosity than at − 0.2 V and − 0.3 V (**Figure 2A**). SEM imaging was used to further compare the nano- and microstructures of the electrodes. Planar control electrodes had a flat surface (**Figure 2B**), whereas electrodes electroplated at 0 V displayed a substantial number of evenly dispersed globular nuclei (**Figure 2C**). At −0.1 V, larger globular nuclei up to 300 nm formed with clearly distinguishable grain boundaries (**Figure 2D**). As the deposition potential was further reduced to −0.2 V and −0.3 V, scattered secondary nuclei resulted in more ramified and heterogeneous structures with significantly taller platinum deposits at the edges than at the centers (**Figures 2E** and **2F**). At −0.4 V, significant nucleation and growth still occurred, but with smaller, more evenly dispersed dendrites, producing nanoscale pores (**Figure 2G**). The variations in deposit morphology at different electroplating potentials can be explained by a transition from a charge transfer-limited regime at lower potentials to a more mass transport-limited regime with increased overpotential (**Figure 2H**) (18).

**Figure 2.**
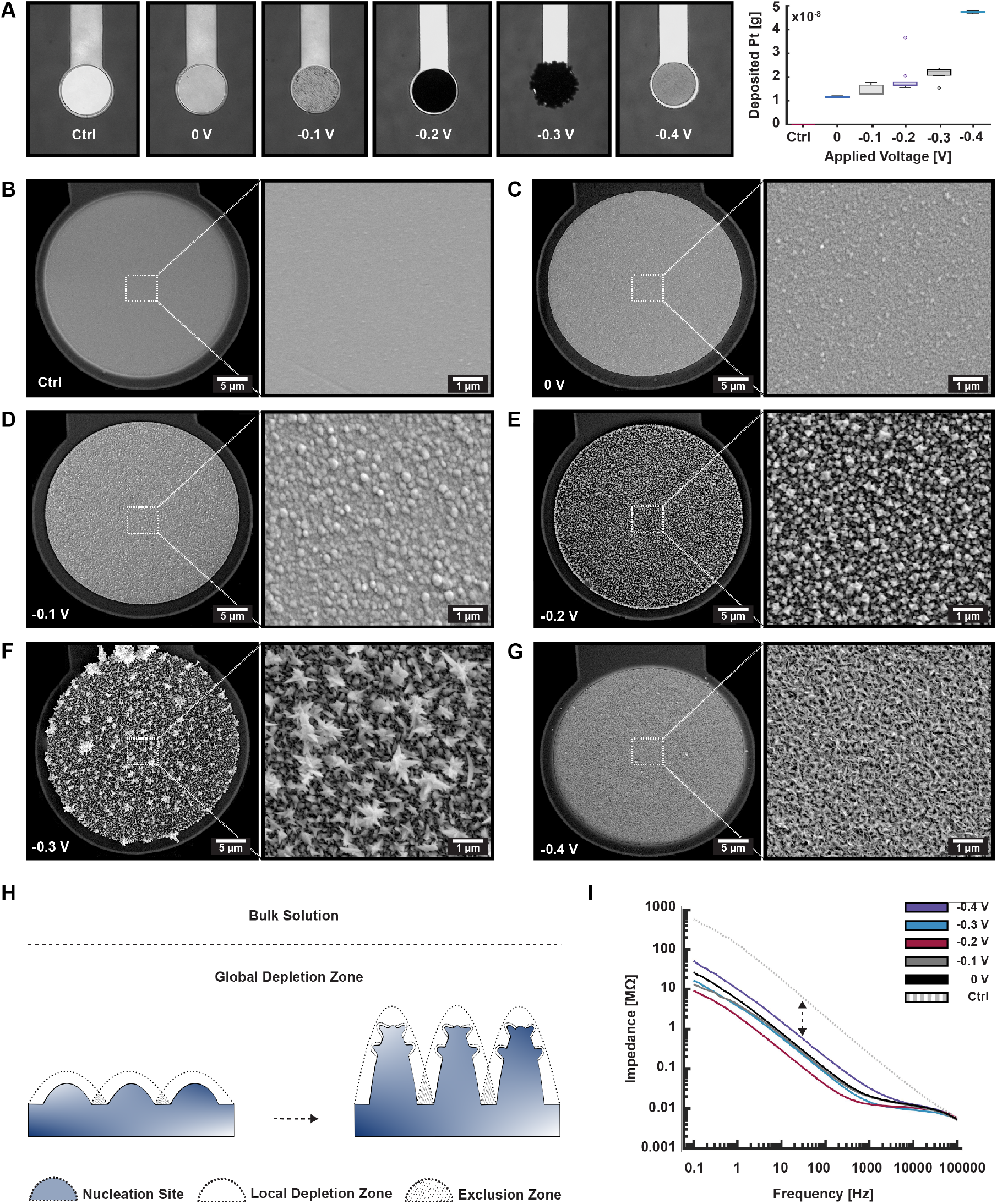
Nano- and microporous platinum electroplated using chronoamperometry. **(A.)** Optical microscopy images showing the increase in porosity, and hence light absorbance, of the electroplated electrodes with decreasing voltage down to −0.3 V. Although the amount of deposited platinum increased with further reduction in deposition potential, the porosity appeared to decrease beyond −0.3 V. **(B.)** Control electrode prior to electrochemical deposition of platinum. **(C.)** - **(G.)** SEM micrographs of 30 µm diameter electrodes electroplated at 0 V, −0.1 V, −0.2 V, −0.3 V and −0.4 V, respectively. The micrographs reveal that electroplating at −0.2 V and −0.3 V produced the most porous electrodes with noticeable edge effects. Electroplating at −0.4 V resulted in evenly layered, highly nanoporous structures. **(H.)** Illustration depicting the mass transport limited nucleation and growth of platinum on the microelectrodes, with local and global depletion zones forming around the nucleation sites, promoting high porosity. **(I.)** Electrochemical Impedance Spectroscopy (EIS) measurements confirmed that increased electrode porosity, due to a higher electroactive surface area, resulted in lowered impedance.

To characterize the impact of the formed structures on the electrodes' impedance, EIS measurements were conducted in PBS solution over a range from 100 kHz to −0.1 Hz. All the nano- and microporous electrodes exhibited impedances' one to two orders of magnitude lower than the control electrodes (**Figure 2I**). At the lowest frequency, 0.1 Hz, the electrodes electroplated at −0.2 V and −0.3 V performed the best, with 63 and 35 times lower impedances compared to the planar controls, respectively. Among the porous electrodes, those deposited at 0.4 V had the highest impedance, yet still 11 times lower than the control electrodes. At higher frequencies, where spikes are typically detected (300 Hz to 3000 Hz), the electrodes electroplated at −0.3 V had the lowest overall impedance, with at least 9.7 times lower impedance than the controls. The electrodes electroplated at −0.4 V had at least 5.3 times lower impedance than the controls in this frequency range. At 1000 Hz, a highly relevant frequency band for spiking neural activity, the electrodes deposited at 0 V, ™0.1 V, −0.2 V, −0.3 V, and −0.4 V, as well as the controls, had impedances of 0.021 MΩ, 0.020 MΩ, 0.013 MΩ, 0.015 MΩ, 0.032 MΩ, and 0.24 MΩ, respectively. Combined with the imaging results, these findings clearly show that all the porous electrodes had a significantly larger electroactive surface area compared to the controls, leading to substantially reduced impedances.

### Chronoamperometric Depositions Yield Consistent Porosity Irrespective of the Electrode Size

An important consideration when designing microelectrodes for neuronal recordings is the electrode size. Larger electrodes increase the likelihood of neurons positioning themselves on or near the electrodes, while smaller electrodes provide higher selectivity by recording from fewer neurons. Establishing a reliable fabrication protocol is crucial for ensuring that the electrochemical deposition process creates porous structures with consistent thickness and without discernible edge effects, regardless of the working electrode size. In this study, different deposition parameters were tested on electrodes of various sizes relevant for electrophysiological recordings, ranging from 10 µm to 100 µm. When chronoamperometric depositions were performed at −0.4 V, evenly layered nanoporous platinum was formed regardless of electrode size (**Figures 3A** - **3D**). The electroplated structures were also consistent and independent of electrode size for depositions at −0.2 V and −0.3 V, producing the most microporous electrodes (**Figure S2**). However, compared to electrodes plated at −0.4 V, these electrodes exhibited more distinct edge effects, particularly for the smallest electrodes between 10 µm to 30 µm in diameter. In contrast, chronopotentiometric depositions resulted in electrode porosity that was clearly dependent on electrode size (**Figures 3E** - **3H**). This can be attributed to the continuously increasing surface area over time, which disperses the applied constant current over a larger total surface area. This made smaller electrodes (10 µm to 30 µm in diameter) more prone to extensive platinum growth of along their edges compared to their centers (**Figures 3E** and **3F**). These results demonstrate that chronoamperometry facilitates more reproducible and consistent electroplating independent of electrode size compared to chronopotentiometry.

**Figure 3.**
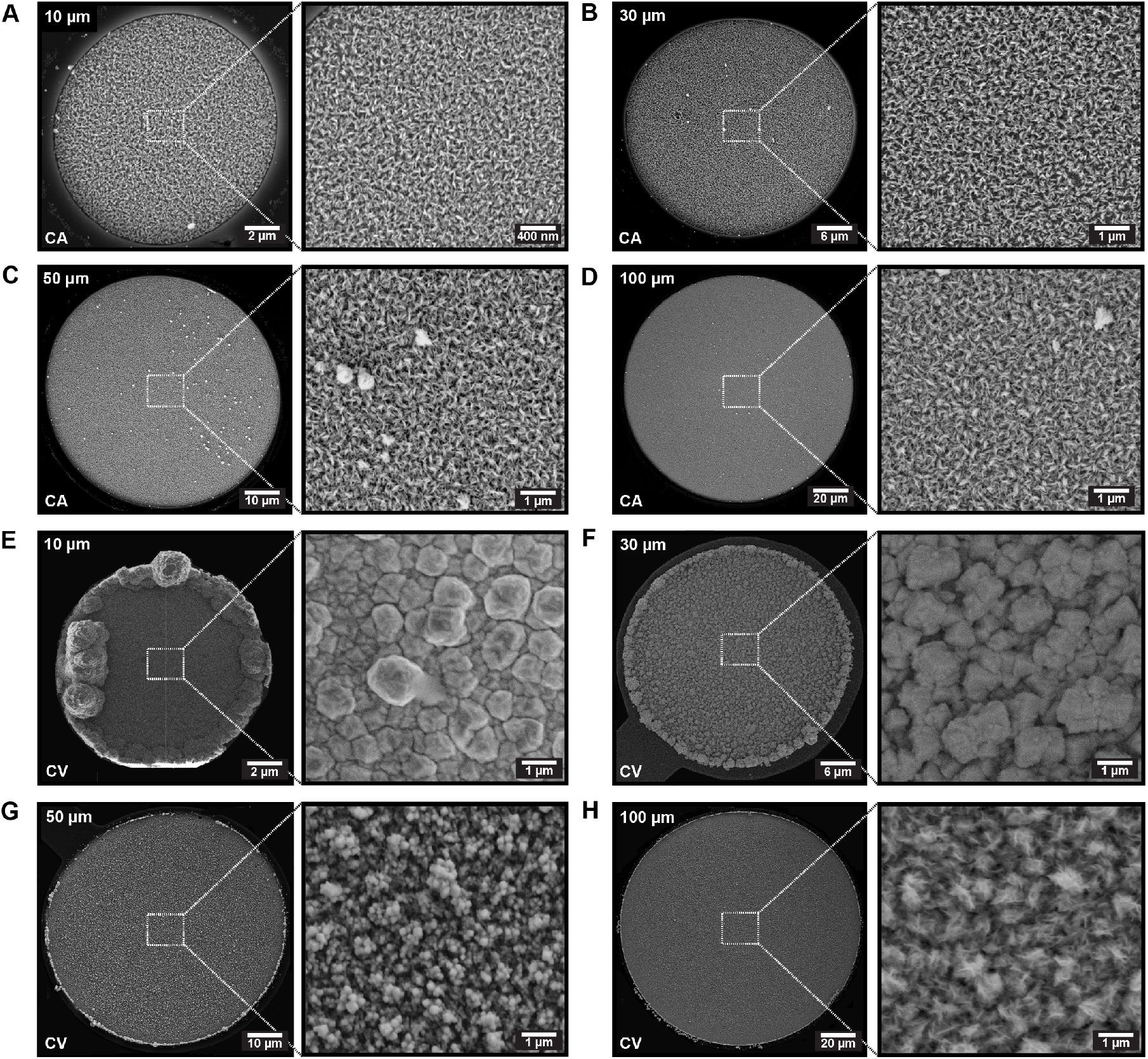
Dependence of electrode size on electrode porosity when applying chronoamperometric and chronopotentiometric depositions. **(A.)** - **(D.)** Micrographs demonstrating that electrodes with diameters ranging from 10 µm to 100 µm exhibited evenly layered, nanoporous platinum deposits without discernible edge effects, irrespective of electrode size, when performing chronoamperometric depositions at −0.4 V for 3 min. **(E.)** - **(F.)** Micrographs illustrating the gradual transition into smaller and more evenly dispersed dendrites as the electrode size increased from 10 µm to 100 µm when applying chronopotentiometric depositions at 0.1 nA/µm for 5 min.

To assess the impact of electrode size on performance in electrophysiological recordings of neurons, cells were plated on MEAs with electrodes electroplated at −0.4 V ranging from 10 µm to 100 µm in diameter, randomly distributed across the interfaces. Recordings conducted during early maturation at 14 DIV indicated a clear dependence of electrode size on the percentage of active electrodes, i.e., electrodes showing strong coupling with one or several neurons (**Figure 4A** and **Table SI**). Electrodes with a 100 µm diameter had over 90% active electrodes, while those 50 µm or smaller had less than 60% active electrodes at this time point. By 26 DIV, when the networks had matured, electrodes 30 µm or larger approached 100% active electrodes, whereas electrodes of 10 µm and 20 µm in diameter had above 60% active electrodes. The SNR followed a similar trend, with significantly higher SNR for larger electrodes at 14 DIV, and a non-significant but higher SNR at 26 DIV (**Figure 4B**). Conversely, the median spike amplitude and variance in the noise band showed higher amplitudes and lower noise variance for smaller electrodes (**Figures 4C** and **4D**, respectively). These findings underscore the importance of balancing selectivity and electrochemical performance when selecting electrode size.

**Figure 4.**
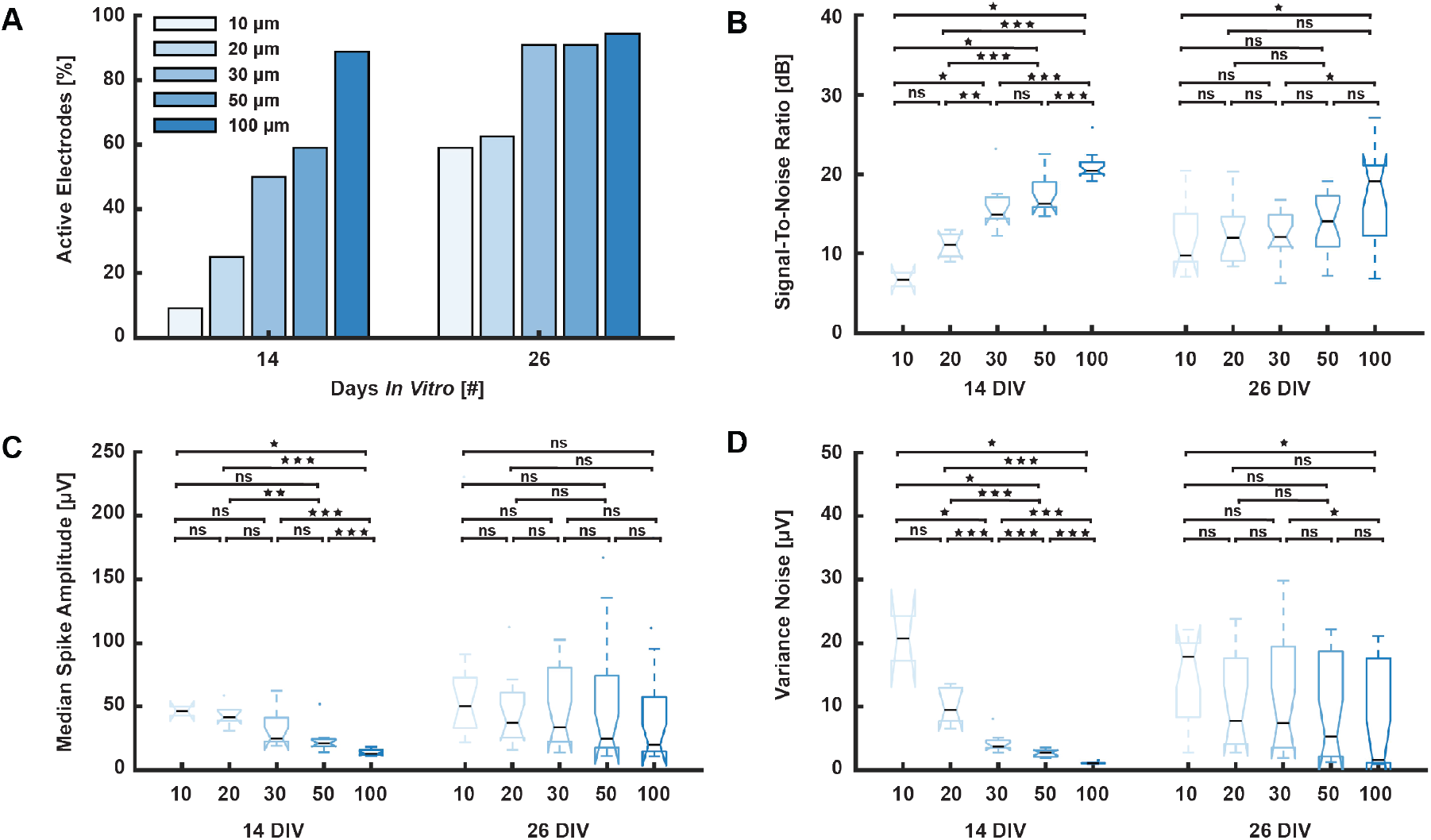
Impact of electrode size on electrophysiological performance of porous platinum electrodes. **(A.)** At 14 days *in vitro* (DIV), the electrode size significantly influenced the percentage of electrodes detecting neuronal activity. Less than 10% of the 10 µm diameter electrodes were able to detect activity, while more than 90% of the 100 µm diameter electrodes were active. By 26 DIV, over 60% of electrodes of all sizes were active, with more than 95% of electrodes 30 µm in diameter or larger detecting activity. **(B.)** At 14 DIV, the Signal-To-Noise (SNR) ratio showed a clear dependence on electrode size, with 100 µm diameter electrodes having three times higher SNR than 10 µm electrodes. At 26 DIV, the differences in SNR between electrode sizes were less pronounced. **(C.)** The median spike amplitude displayed an opposite trend, with 10 µm diameter electrodes showing the highest amplitudes at 14 DIV. By 26 DIV, no significant differences in spike amplitudes were observed across electrode sizes. **(D.)** The variance in the noise band was significantly lower for the largest electrodes at 14 DIV, but the differences across electrode sizes were less pronounced at 26 DIV.

### Chronoamperometric Depositions Yield Consistent Porosity Irrespective of Deposition Time

In addition to the applied voltage or current amplitude, deposition time is a crucial parameter in electrochemical depositions. To evaluate its impact on the final nano- and microstructures, chronoamperometric and chronopotentiometric depositions were conducted for varying durations. Chronoamperometric depositions at −0.4 V were performed for periods ranging from 2 min to 10 min. These depositions produced evenly layered, nanoporous structures without discernible edge effects on the electrodes, irrespective of the deposition time (**Figures 5A** and **5B**). Conversely, chronopotentiometric depositions resulted in larger, globular structures as the deposition time increased (**Figures 5C** - **5F**). After 30 min of electroplating, the initial nuclei had merged into a homogeneously layered surface with distinguishable grain boundaries (**Figure 5F**). Additionally, edge effects became significantly more prevalent with longer deposition times for these electrodes. These findings underscore the versatility of chronoamperometric depositions, as the process yields the same electrode porosity independent of deposition time and electrode size.

**Figure 5.**
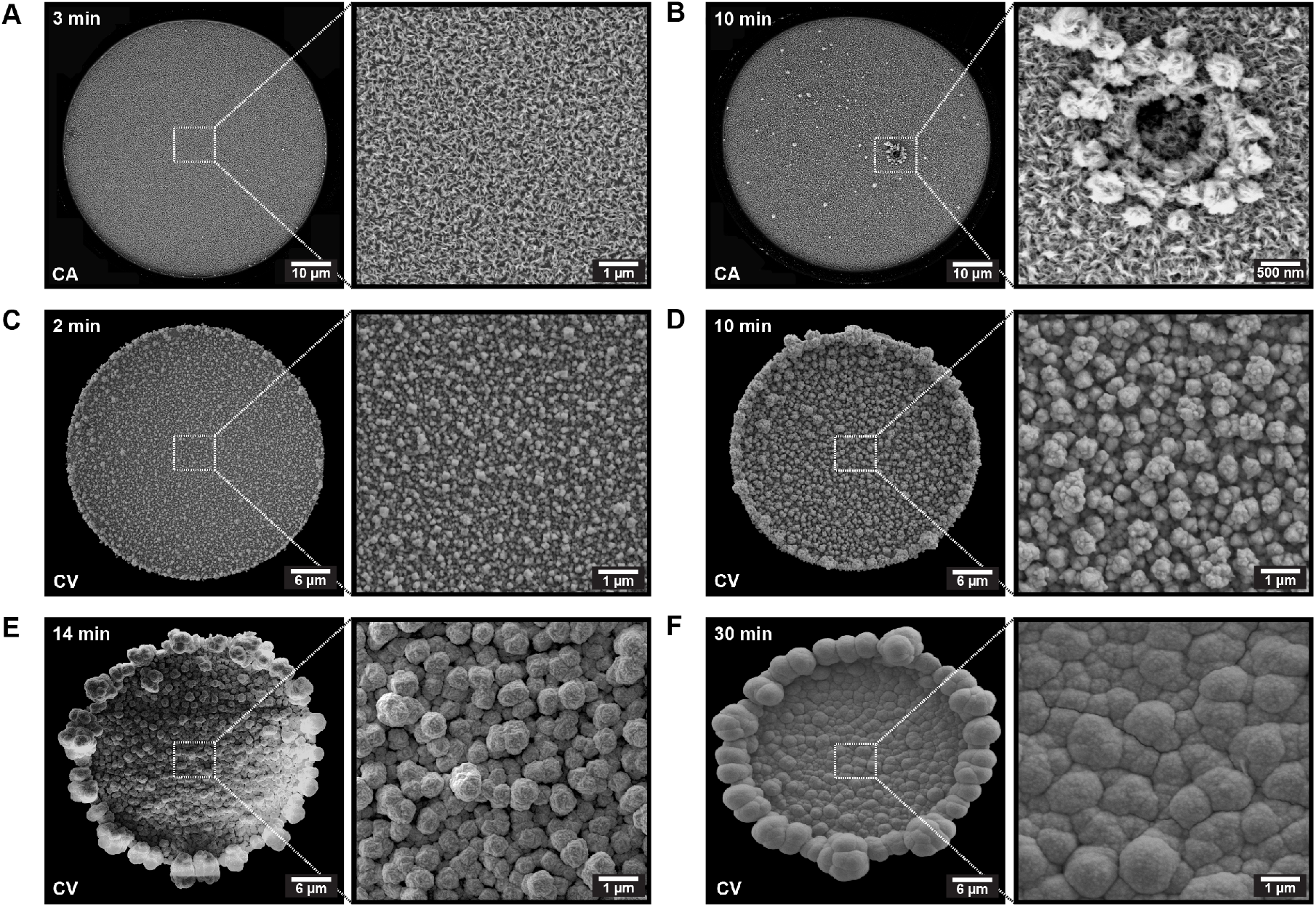
Dependence of deposition time on the electrode porosity for chronoamperometric and chronopotentiometric electrodepositions. **(A.)** - **(B.)** Chronoam- perometric depositions at −0.4 V resulted in evenly layered, nanoporous platinum structures, irrespective of the deposition time. The defect in the electrode after a 10 min deposition reveals clearly defined layers of nanoporous platinum throughout the cross-section of the electrode. **(C.)** - **(F.)** Chronopotentiometric depositions showed a growth of nuclei into larger, globular structures as the deposition time extended beyond 2 min, eventually merging into a homogeneous layer after 30 min.

### Addition of Formic Acid and Application of Dynamic Deposition Schemes Decrease Reproducibility of the Electroplating

Additives such as lead acetate or formic acid are commonly added to the electrolyte bath to facilitate nucleation and improve the adhesion of the porous platinum deposits (10, 35). To evaluate the impact of additives in the electrolyte bath, formic acid (HCOOH) was included in the electrolyte solution prior to performing chronoamperometric depositions between 0 V to − 0.4 V. While the addition of HCOOH promoted the growth of highly porous platinum structures, it also led to significant edge effects at all deposition potentials (**Figure S3**). To mitigate edge effects and reduce the formation of heterogeneous and ramified structures, various dynamic deposition protocols were tested, including linearly decreasing voltages, cyclic voltammetry, and pulsed depositions. These depositions were performed both with and without the addition of formic acid to assess their impact on the resulting nano- and microstructures (**Figures S4** and **S5**). Although these methods produced a variety of electrode morphologies, they did not significantly reduce edge effects or the presence of heterogeneous microstructures. Consequently, the reproducibility of these approaches was lower than that of chronoamperometry, leading to their exclusion from further electrochemical and electrophysiological performance testing.

### Heterogeneous, Microporous Electrodes Yield Lower Noise Bands, While Evenly Layered, Nanoporous Electrodes Yield Higher Spike Amplitudes

To evaluate the effect of electrode porosity on the data quality obtained from electrophysiological recordings, MEAs were designed to include a mixture of planar control electrodes and electrodes electroplated at either −0.3 V or −0.4 V. 6 MEAs were used for these experiments, each containing 20 electrodes of the different types. By including different electrode types on the same MEAs, potential biases from differences between distinct neuronal networks were avoided. The chosen deposition parameters allowed for a comparison between the most microporous electrodes, having the lowest impedance, and the evenly layered, nanoporous electrodes. Microscopic examination showed no apparent differences in biocompatibility across the electrode types, with prominent neuronal growth observed on all platforms (**Figure 6A**). Electron microscopy further revealed that even the most porous electrodes supported significant neurite meshes extending across their surfaces (**Figure 6B**).

**Figure 6.**
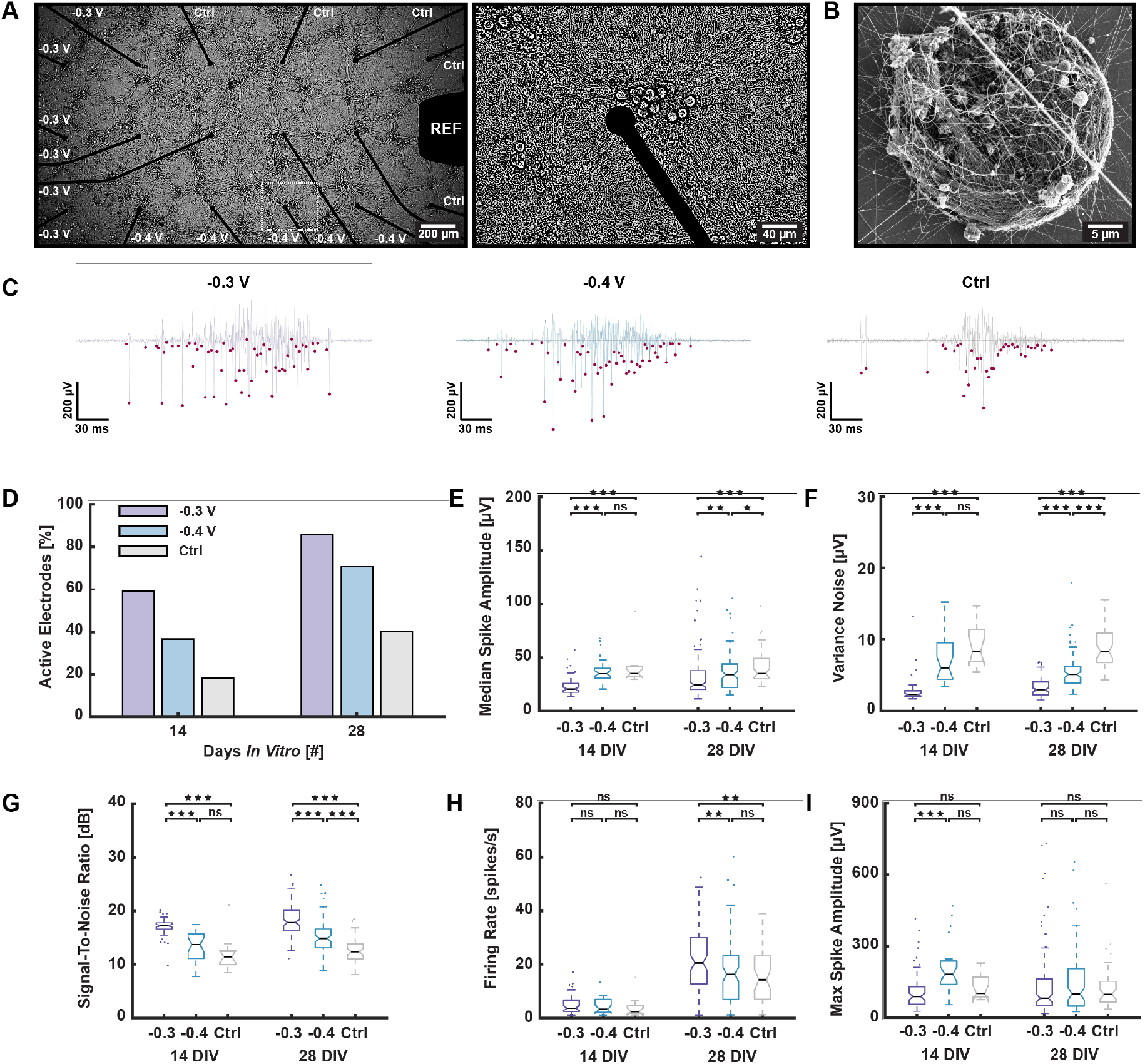
Electrophysiological performance and biocompatibility of porous platinum electrodes in neural cell cultures. **(A.)** Micrographs showing the high biocompatibility of the microelectrodes independent of porosity, with extensive neuronal processes across all electrodes. **(B.)** Scanning electron microscopy (SEM) image showing extensive neurite branching throughout a highly porous electrode electroplated at −0.3 V. **(C.)** Representative voltage traces demonstrating high signal-to-noise (SNR) ratios for electrodes electroplated at −0.3 V, −0.4 V, and planar controls, respectively. Pink specks represent spikes detected using the precise timing spike detection (PTSD) algorithm. **(D.)** A significantly higher proportion of the nano- and microporous electrodes were active throughout the experimental period compared to planar control electrodes. **(E.)** - **(F.)** Both median spike amplitudes and variance in the noise bands were significantly lower for electrodes electroplated at −0.3 V compared to electrodes electroplated at −0.4 V and planar controls. **(G.)** The SNR followed an opposite trend, with the highest SNR for the most microporous electrodes and the lowest SNR for the planar controls. **(H.)** Firing rates, indicating the sensitivity of the electrodes, were comparably high for all electrode types at 14 days *in vitro* (DIV), but were significantly higher for the most microporous electrodes at 28 DIV. **(I.)** The maximum spike amplitudes were significantly higher for the nanoporous electrodes electroplated at −0.4 V at 14 DIV, and non-significantly higher at 28 DIV.

Common methods for assessing biocompatibility, such as live/dead assays, are prone to inaccuracies due to cell density variations and the detachment of dead cells. Therefore, biocompatibility was indirectly assessed through electrophysiological recordings, providing a more reliable and dynamic evaluation of neuronal health and network activity, as well as insights into cell adhesion, neuron-electrode coupling, and the potential impact of edge effects on cell proximity to the electrodes. All electrode types exhibited prominent activity, with spike amplitudes ranging from 20 µV to over 300 µV at both 14 and 28 DIV (**Figure 6C**), consistent with typical extracellular action potentials and indicative of a high degree of biocompatibility (3). When comparing the electrophysiological activity of the cells at 14 and 28 DIV, a higher percentage of electrodes electroplated at −0.3 V were active compared to electrodes electroplated at −0.4 V and planar controls (**Figure 6D** and **Table SII**). The most porous electrodes showed significantly lower median spike amplitudes but also lower variance in the noise band compared to the other electrodes (**Figures 6E** - **6F**). Despite the lower median spike amplitudes, the low noise contributed to a significantly higher SNR for these electrodes (**Figure 6G**). The difference in firing rate detected by the three electrode types, indicating electrode sensitivity, was non-significant at 14 DIV but significantly higher for the electrodes electroplated at 0.3 V at 28 DIV compared to the electrodes electroplated at −0.4 V and the planar controls (**Figure 6H**). The maximum spike amplitudes detected by each electrode were, on average, significantly higher for both the electrodes electroplated at −0.4 V and the planar controls compared to the more porous electrodes electroplated at −0.3 V (**Figure 6I**). While the most microporous electrodes had significantly higher SNR, the higher spike amplitudes of the evenly layered, nanoporous electrodes indicate that the neurons may have had better positioning in relation to the electrodes, contributing to better signal acquisition.

It is worth noting that several factors, in addition to electrode morphology, can influence the signal quality of the electrodes. For instance, planar electrodes were found to be more prone to astrocyte coverage, which can shiel the electrodes from the neurons (**Figure S6A**). Scanning electron microscopy revealed astrocytes covering the electrodes, potentially undergoing astrogliosis (**Figure S6B**) (36–38). Another issue was the high fasciculation and formation of thick axonal bundles, which detached from the substrate as the networks matured, leaving a significant gap between the neurons and the electrodes (**Figure S6C**). These results highlight the importance of evaluating biocompatibility and cell-electrode interactions when analyzing electrode performance, in addition to their electrochemical properties.

## Discussion

As shown in both this and previous studies, the electrodeposition of nanoporous platinum is a versatile process that can be fine-tuned to create a variety of nano- and microstructures (35, 39). Numerous parameters influence the characteristics of the formed deposits, including the choice of additives, working electrode material, temperature of the electrochemical bath, deposition current/voltage, and deposition time (18, 19, 40). Although the exact mechanisms behind the nucleation and growth of such structures are not fully understood, our findings support the work of Boehler *et al*., which suggest that electroplating without additives promotes a more predictable and reproducible deposition process (12). Additionally, our results indicate that chronoamperometric depositions produce more consistent structures regardless of electrode size and deposition time. While low overpotentials resulted in globular structures connected by grain boundaries, higher overpotentials led to more dispersed secondary nucleation from depletion zones forming around the primary nuclei, consistent with previous studies (18). By fine-tuning the overpotential to −0.4 V, we achieved highly nanoporous structures without discernible edge effects on hundreds of electrodes in parallel.

When designing microelectrodes for biosensor applications, it’s crucial to consider their biocompatibility (41, 42). While many studies assess biocompatibility solely through cell viability, the impact of electrode morphology on neuron positioning relative to the electrodes is often overlooked. Although higher porosity typically improves electrochemical performance, it can lead to significant edge effects and rough electrode morphologies, affecting interactions with neurons and glial cells. Moreover, these protruding structures are fragile and may break off during cell culturing, potentially being engulfed by cells. Our study found that the most microporous electrodes, electroplated at −0.3 V, exhibited higher activity and signal-to-noise ratio (SNR) compared to more evenly layered nanoporous electrodes at −0.4 V and planar controls. This higher SNR was primarily attributed to lower noise variance, as the less porous electrodes detected higher median and maximum spike amplitudes. Theoretically, the highest spike amplitudes are detected at the perisomatic area, and the amplitude will primarily be determined by the proximity between the neuron and the electrode (3). While the most porous electrodes may better detect low-amplitude signals from neurons farther away, their tall edges and large protrusions may increase the distance between neurons and electrodes. Conversely, less microporous electrodes may not detect signals from distant cells but may promote neuron positioning closer to or on the electrodes. This may contribute to subsampling due to fewer active electrodes (43), but it could also simplify spike sorting as electrodes are more selective to individual neurons (44). Therefore, while microelectrodes with large edge effects exhibit high SNR, using homogeneously layered nanoporous electrodes may generally be advantageous.

The current study has focused on the application of nanoporous platinum for extracellular electrodes used with *in vitro* engineered neural networks. However, as demonstrated, the established deposition protocol is highly versatile, accommodating different electrode sizes, electrode numbers, and deposition times, making it well-suited for a variety of applications, both *in vitro* and *in vivo*. Nanoporous platinum is also being explored for use in *in vivo* implantable electrodes, where it is crucial to prevent the breakage of nano- and microporous structures, as fragments could lead to cytotoxicity, tissue damage, or cellular engulfment (12, 21, 45, 46). Moving forward, it will be important to evaluate the long-term biological responses in *in vivo* applications, such as glial scarring, inflammation, and neural network stability, to fully understand the chronic effects of these electrodes in living tissues.

## Conclusion

Nano- and microporous electrodes offer enhanced signal-to-noise ratio, making them appealing for electrophysiological applications. In this study, we manipulated the electrochemical deposition of platinum to generate varied electrode morphologies at nano- and micrometer scales. While higher porosity improved electrochemical electrode performance, rougher morphology negatively affected neuron-electrode coupling, likely due to prominent edge effects. Evenly layered, nanoporous electrodes exhibited higher median and maximum spike amplitudes, suggesting closer proximity to perisomatic neuronal areas. These results highlight the importance of considering not only electrochemical characteristics and cell viability when assessing electrode performance, but also the impact of the morphology on the neuron-electrode junction. In conclusion, we have demonstrated a highly versatile fabrication protocol for creating evenly layered, nanoporous platinum microelectrodes with an optimal trade-off between biocompatibility, electrochemical, and electrophysiological performance.

## AUTHOR CONTRIBUTIONS

The author contributions follow the CRediT system. **NWH**: Conceptualization, Methodology, Software, Investigation (chip design & manufacturing, cell experiments, electrophysiology, SEM and bioSEM imaging, formal analysis), Writing – Original Draft, Visualization. **LI**: Methodology, Investigation (chip manufacturing, SEM imaging), Writing – Review & Editing. **PAK**: Methodology, Writing – Review & Editing. **AS, IS**: Conceptualization, Writing – Review & Editing, Resources, Funding Acquisition. **PS**: Conceptualization, Methodology, Writing – Review & Editing, Resources, Funding Acquisition.

## ACKNOWLEDGEMENTS

We would like to thank engineers Mark Chiappa, Martijn de Roosz and Mathilde Barriet at NTNU NanoLab for technical assistance. Furthermore, we acknowledge Nan Tostrup Skogaker for training and access to the electron microscopy core facility, NTNU. Prof. Michela Chiappalone and Prof. Sergio Martinoia, University of Genova are acknowledged for generously providing the scripts for the Precise Timing Spike Detection algorithm. We would also like to thank professor Svein Sunde for insights and valuable discussions on the electrochemical characterization.

## DECLARATION OF GENERATIVE AI AND AI-ASSISTED TECHNOLOGIES IN THE WRITING PROCESS

During the preparation of this work the authors used ChatGPT in order to improve language and readability. After using this tool, the authors reviewed and edited the content as needed and take full responsibility for the content of the publication.

