## Supplementary Materials for "Nanoporous Platinum Microelectrode Arrays for Neuroscience Applications"

**Nicolai Winter-Hjelm[1,✉], Leik Isdal[2], Peter A. Köllensperger[3], Axel Sandvig[1,4], Ioanna Sandvig[1], and Pawel Sikorski[2,✉]**

[1]Department of Neuromedicine and Movement Science, Faculty of Medicine and Health Sciences, Norwegian University of Science and Technology (NTNU), Norway
[2]Department of Physics, Faculty of Natural Sciences, Norwegian University of Science and Technology (NTNU), Trondheim, Norway
[3]NTNU NanoLab, Norwegian University of Science and Technology (NTNU), Trondheim, Norway
[4]Department of Neurology and Clinical Neurophysiology, St Olav´s University Hospital, Trondheim, Norway

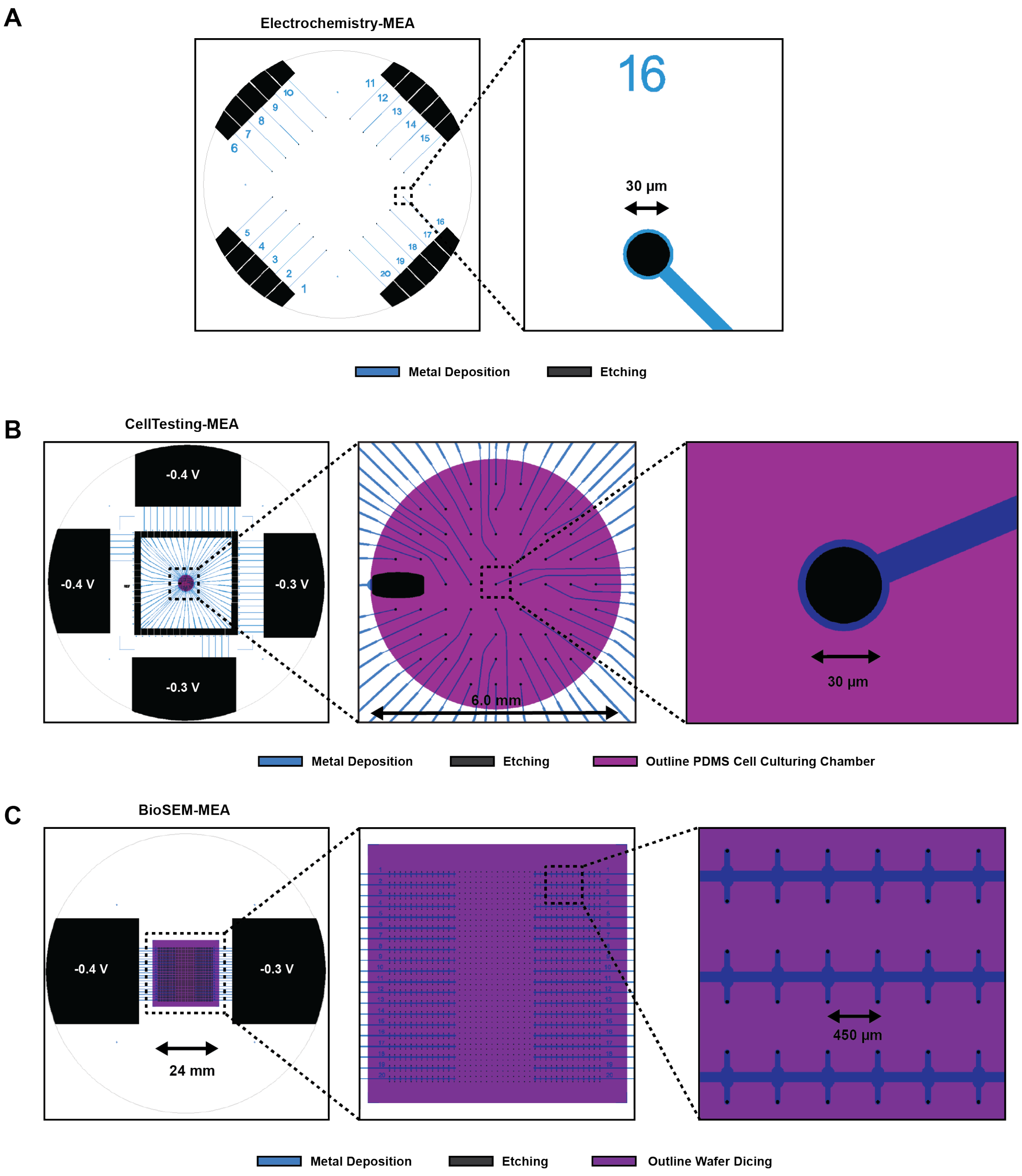


**Figure S1** | **CAD designs of the microelectrode arrays**. **(A.)** Design utilized for electrochemical testing and scanning electron microscopy of the microelectrodes with varying deposition parameters. Layer 1 represents the metallization design, forming the microelectrodes and corresponding contact pads. Layer 2 serves as the etch mask for etching through the passivation layer. Each microelectrode measures $30\,\mu\mathrm{m}$ in diameter and connects to individual contact pads at the periphery of the 4" wafer to support distinct electrochemical depositions. **(B.)** Design employed for electrophysiology of neural cell cultures growing on the microelectrode arrays. The four large contact pads at the wafer periphery connects the MEA to the potentiostat during electroplating. After deposition, the wafer is cut into a 49 x 49 mm square using a wafer saw along the four corner marks, disconnecting the 60 smaller contact pads from the larger ones. The third layer, *outline PDMS cell culturing chamber*, outlines the PDMS chambers. **(C.)** Design utilized for scanning electron microscopy (SEM) imaging of neurons growing atop the microelectrodes. 520 electrodes were electroplated at $-0.4\,\mathrm{V}$, 520 electrodes at $-0.3\,\mathrm{V}$, and 520 were used as planar controls. As the electrodes were not employed for electrophysiology, they were not wired to individual contact pads.

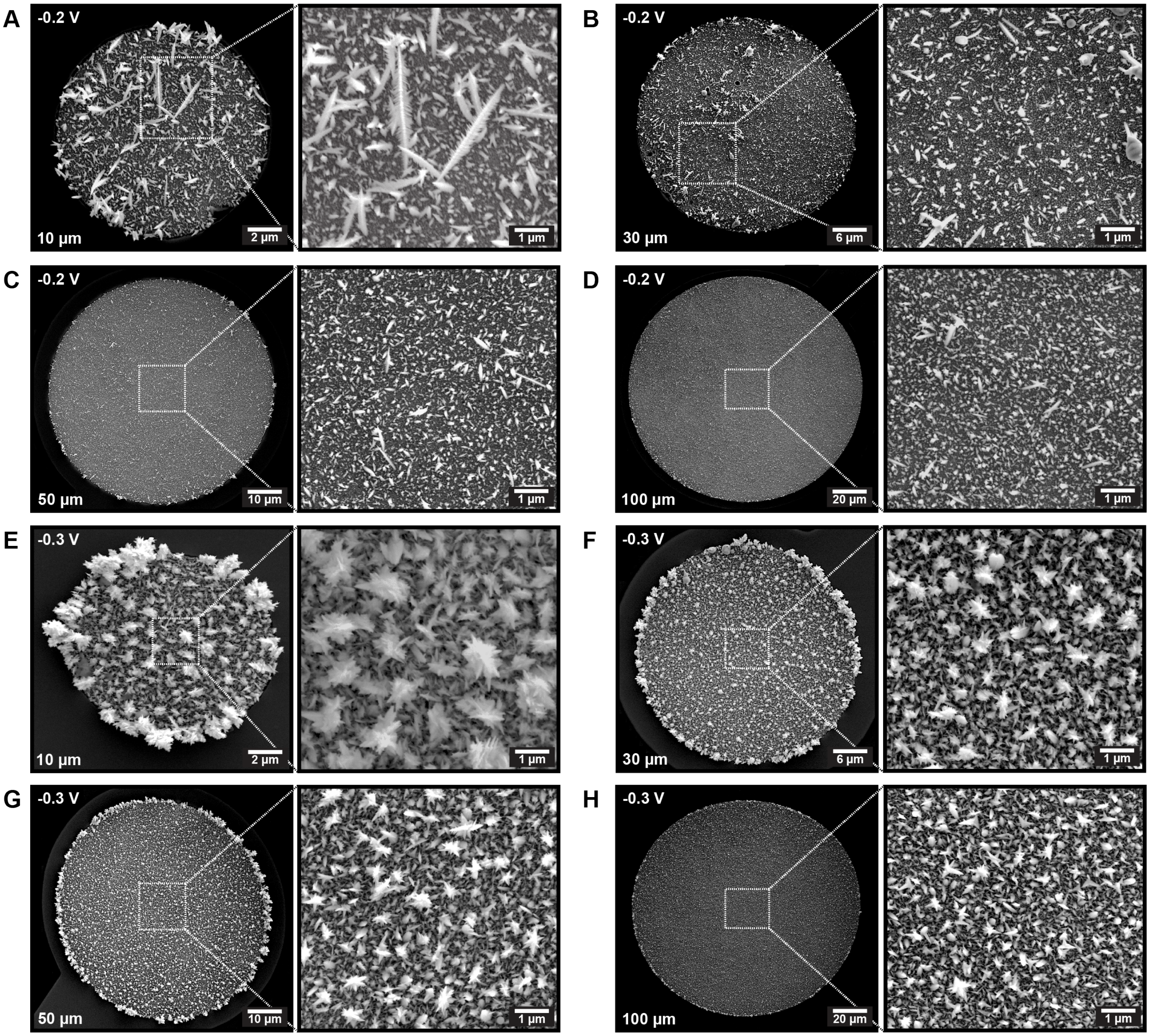


**Figure S2** | **Dependence of electrode size on electrode porosity for chronoamperometric electrodepositions**. **(A.)** - **(D.)** Micrographs depicting electrodes with diameters ranging from $10\,\mu m$ to $100\,\mu m$, exhibiting porous platinum deposits with similar morphology irrespective of electrode size. These deposits were achieved through electroplating using chronoamperometry at $-0.2\,V$ for $3\,min$. **(E.)** - **(H.)** Micrographs illustrating electrodes with diameters ranging from $10\,\mu m$ to $100\,\mu m$, displaying comparable porous platinum deposit morphologies regardless of electrode size. These deposits were obtained through electroplating using chronoamperometry at $-0.3\,V$ for $3\,min$.

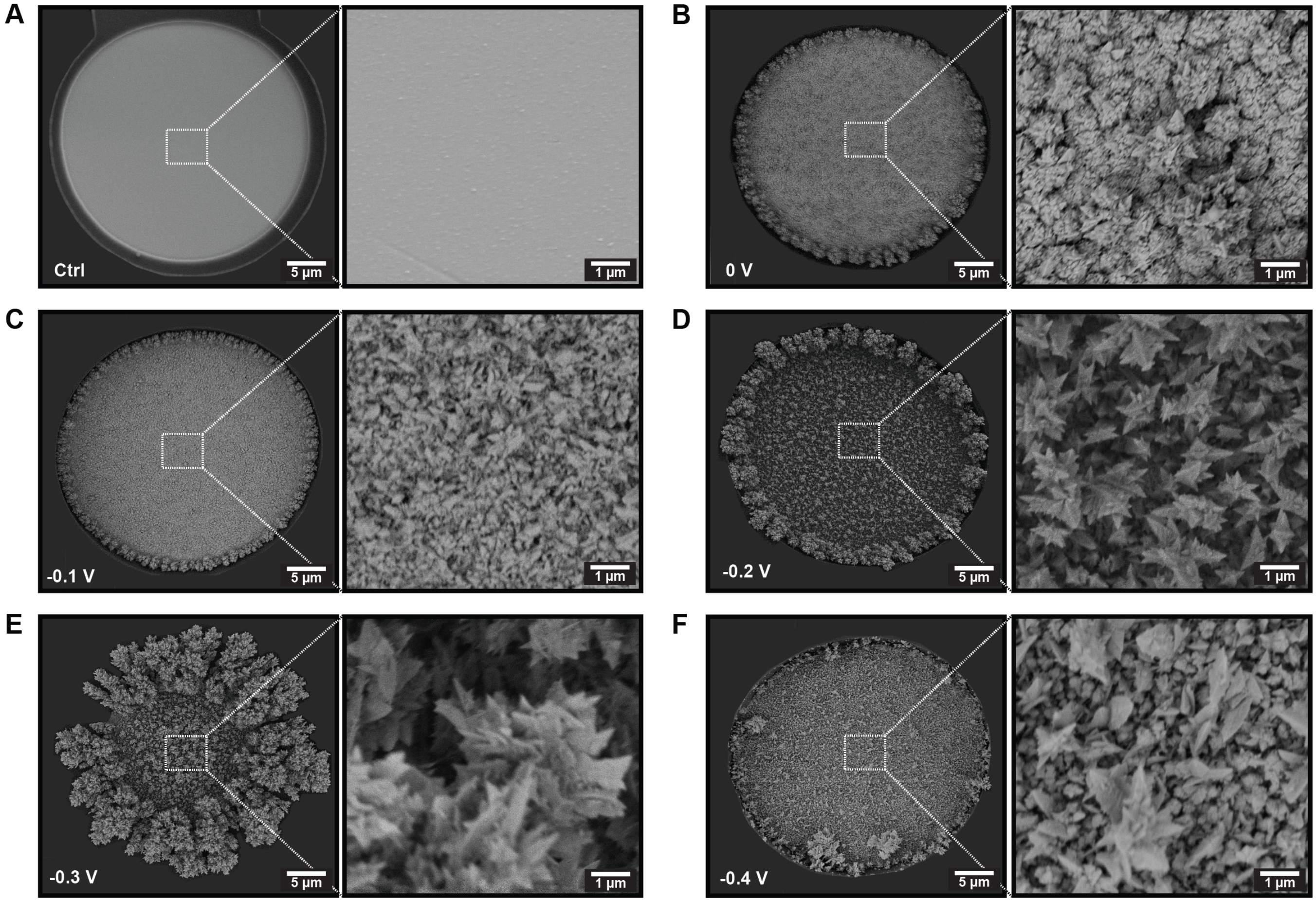


**Figure S3** | **Electrode porosity when using formic acid as an additive during chronoamperometric depositions**. **(A.)** Scanning electron microscopy (SEM) image depicting a planar control electrode before electrochemical deposition of porous platinum. **(B.)** - **(E.)** SEM images demonstrating the gradual increase in porosity with decreased deposition potentials from $0\,\mathrm{V}$ to $-0.3\,\mathrm{V}$, respectively. All depositions lasted for $2\,\mathrm{min}$, and formic acid (HCOOH) served as an additive. **(F.)** SEM image indicating that for chronoamperometric depositions at $-0.4\,\mathrm{V}$, porosity was notably lower than at $-0.3\,\mathrm{V}$. However, the porosity remained significantly higher and exhibited more discernible edge effects compared to when applying the same deposition potential without additives.

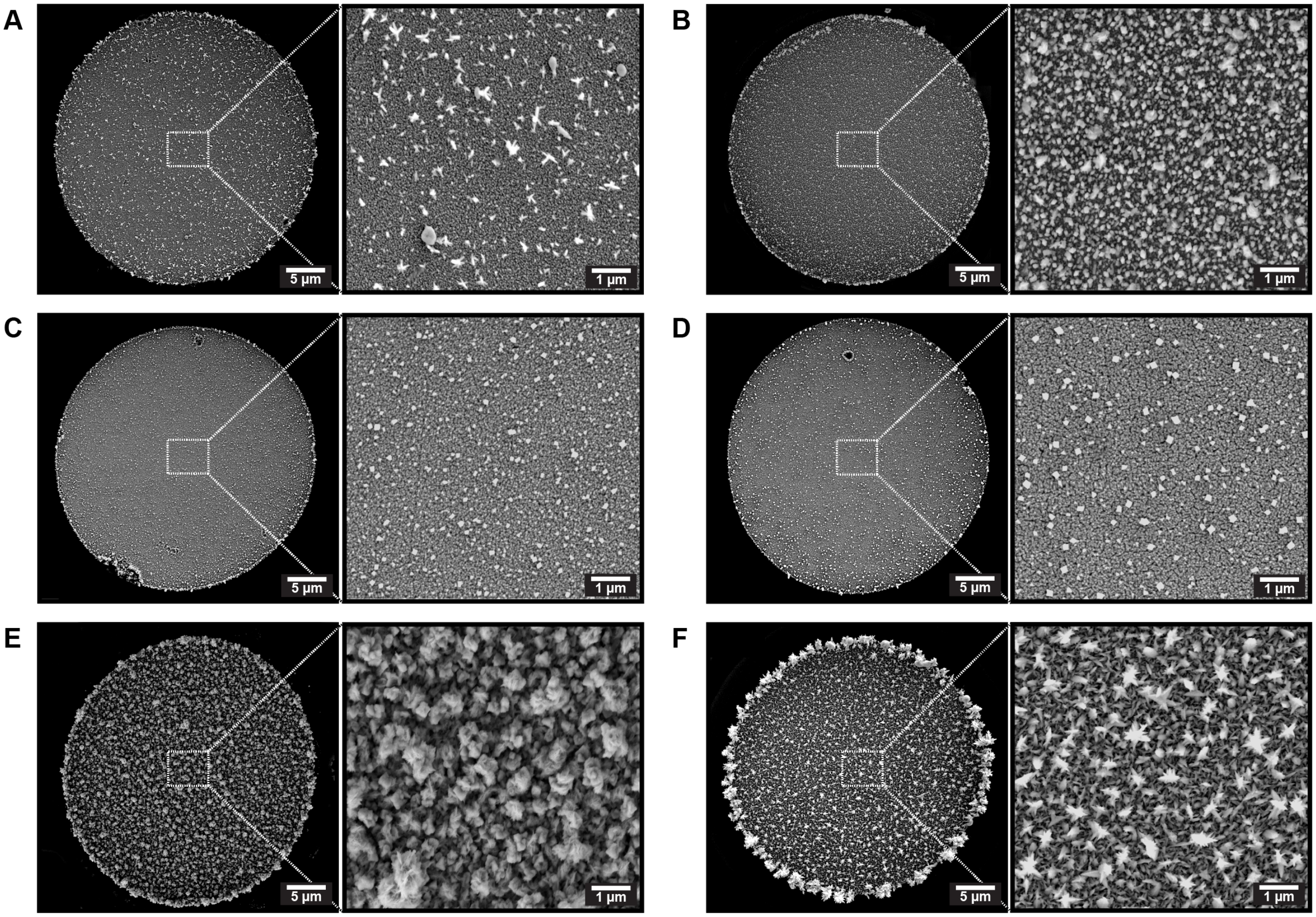


**Figure S4** | **Electrode porosity when applying dynamic deposition schemes**. **(A.)** Linearly decreasing voltage from $0.3\,\mathrm{V}$ to $-0.3\,\mathrm{V}$ at $0.05\,\mathrm{V/s}$. **(B.)** Linearly decreasing voltage from $-0.2\,\mathrm{V}$ to $-0.4\,\mathrm{V}$ at $0.0017\,\mathrm{V/s}$. **(C.)** Cyclic voltammetry for 10 cycles from $0.3\,\mathrm{V}$ to $-0.3\,\mathrm{V}$ at $0.1\,\mathrm{V/s}$. **(D.)** Cyclic voltammetry for 60 cycles from $0.3\,\mathrm{V}$ to $-0.3\,\mathrm{V}$ at $0.6\,\mathrm{V/s}$. **(E.)** Cyclic voltammetry for 10 cycles from $0\,\mathrm{V}$ to $-0.4\,\mathrm{V}$ at $0.1\,\mathrm{V/s}$. **(F.)** Pulsed depositions at $-0.3\,\mathrm{V}$ with $1\,\mathrm{s}$ pulses and $2\,\mathrm{s}$ intervals for $240\,\mathrm{s}$. All electrodes were $30\,\mu\mathrm{m}$ in diameter.

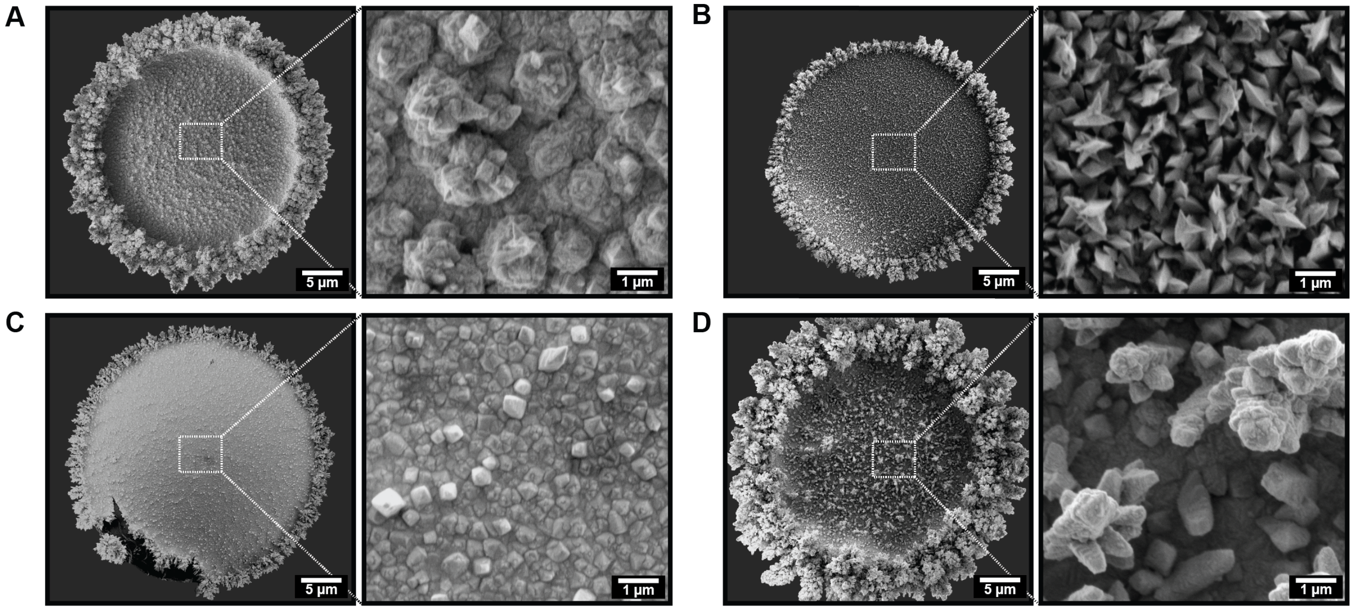


**Figure S5** | **Electrode porosity when using formic acid as an additive, and applying a dynamic deposition scheme**. **(A.)** Cyclic voltammetry for 30 cycles from $0.3\,\mathrm{V}$ to $-0.3\,\mathrm{V}$ at $0.24\,\mathrm{V/s}$. **(B.)** Cyclic voltammetry for 60 cycles from $0.3\,\mathrm{V}$ to $-0.3\,\mathrm{V}$ at $0.36\,\mathrm{V/s}$. **(C.)** Pulsed depositions with $4\,\mathrm{s}$ peaks at $-0.3\,\mathrm{V}$ and $1\,\mathrm{s}$ valleys at $-0.3\,\mathrm{V}$ for $225\,\mathrm{s}$. **(D.)** Pulsed depositions with $1\,\mathrm{s}$ peaks at $-0.3\,\mathrm{V}$ and $4\,\mathrm{s}$ valleys at $-0.3\,\mathrm{V}$ for $225\,\mathrm{s}$. All electrodes were $30\,\mathrm{\mu m}$ in diameter.

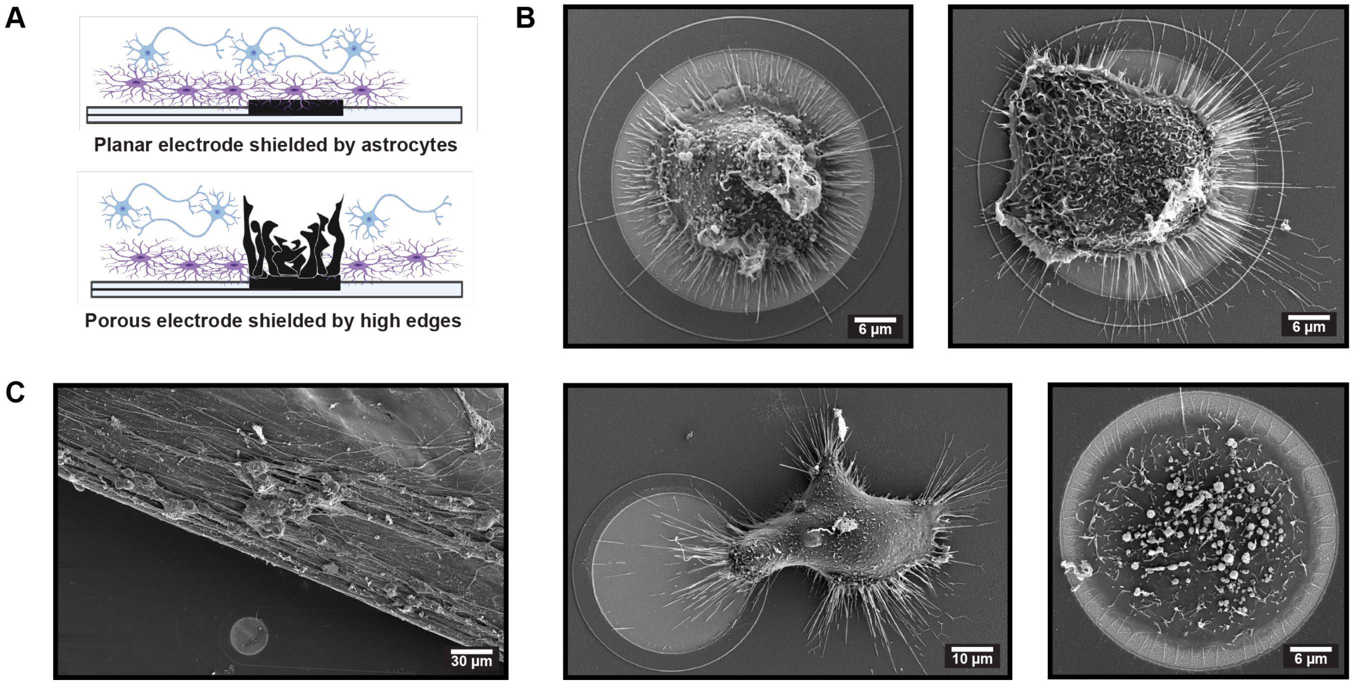


**Figure S6** | **The impact of astrocyte shielding and axonal bundling on neuron-electrode coupling**. **(A.)** Illustrations depicting two possible issues affecting the electrophysiological performance of electrodes: Shielding by astrocytes and shielding of the most porous electrode by their pronounced edge effects and numerous heterogeneous protrusions. **(B.)** Scanning electron microscopy (SEM) images showing electrodes covered with astrocytes, suggesting a selective shielding process indicative of astrogliosis. **(C.)** SEM image illustrating another challenge observed in neural recordings, where neuronal processes detach due to high clustering and bundling, creating a considerable gap between the electrode and the neural tissue.

**Table 1** | Number of active electrodes by electrode size and time point for electrodes electrodeposited at $-0.4\,\mathrm{V}$.

| Time Point [DIV] | MEA [#] | 10 µm | 20 µm | 30 µm | 50 µm | 100 µm |
|---|---|---|---|---|---|---|
| 14 DIV | 1 | 1 / 11 | 3 / 12 | 4 / 11 | 6 / 11 | 7 / 9 |
| | 2 | 1 / 11 | 3 / 12 | 7 / 11 | 7 / 11 | 9 / 9 |
| 26 DIV | 1 | 2 / 11 | 6 / 12 | 9 / 11 | 9 / 11 | 9 / 9 |
| | 2 | 11 / 11 | 9 / 12 | 11 / 11 | 11 / 11 | 8 / 9 |

**Table 2** | Number of active electrodes by electrodeposition voltage and recording time point.

| Time Point [DIV] | MEA [#] | Control | -0.3 V | -0.4 V |
|---|---|---|---|---|
| 14 DIV | 1 | 0 / 19 | 9 / 20 | 3 / 20 |
| | 2 | 0 / 19 | 12 / 20 | 6 / 20 |
| | 3 | 4 / 19 | 12 / 20 | 7 / 20 |
| | 4 | 3 / 19 | 7 / 20 | 6 / 20 |
| | 5 | 5 / 19 | 14 / 20 | 5 / 20 |
| | 6 | 9 / 19 | 12 / 20 | 10 / 20 |
| 28 DIV | 1 | 9 / 19 | 17 / 20 | 17 / 20 |
| | 2 | 6 / 19 | 20 / 20 | 10 / 20 |
| | 3 | 6 / 19 | 16 / 20 | 15 / 20 |
| | 4 | 7 / 19 | 19 / 20 | 19 / 20 |
| | 5 | 10 / 19 | 20 / 20 | 16 / 20 |
| | 6 | 8 / 19 | 16 / 20 | 15 / 20 |
